# Neural Correlates of human cognitive abilities during sleep

**DOI:** 10.1101/130500

**Authors:** Zhuo Fang, Laura B. Ray, Adrian M. Owen, Stuart M. Fogel

## Abstract

Inter-individual differences in sleep spindles are highly correlated with “Reasoning” abilities (problem solving skills; i.e., the ability to employ logic, identify complex patterns), but not Short Term Memory or Verbal abilities. Simultaneous electroencephalography and functional magnetic resonance imaging (EEG-fMRI) have revealed brain activations time-locked to spindles (e.g., thalamic, paralimbic, and motor cortical areas)–yet the functional significance of inter-individual differences in spindle-related brain activation remains to be investigated. Using EEG-fMRI during sleep, we identified, for the first time, the neural activation patterns time-locked to spindles that are correlated with cognitive abilities. Similar to previous studies, activations time-locked to spindles were observed in thalamocortical circuitry and basal ganglia regions. Importantly, spindle-related activation in a subset of these regions were specifically related to inter-individual differences in Reasoning, but not STM or Verbal abilities. These results may help elucidate the physiological mechanisms which support the function of sleep for the capacity for reasoning.

## INTRODUCTION

The sleep spindle is the only known spontaneous neural oscillation that has been identified as an electrophysiological marker of cognitive abilities and aptitudes, that are typically assessed by intelligence quotient (**IQ**) tests (for review, see Fogel & Smith, 2011). As one of the defining features of Stage 2 non-rapid eye movement (**NREM**) sleep, spindles are traditionally defined as neural oscillations between 11 and 16 Hz (Iber et al., 2007), lasting up to ∼3 sec in duration (Rechtschaffen & Kales 1968). Spindles are remarkably stable from night-to-night, but vary considerably from one individual to another, and have even been suggested to be an “electrophysiological fingerprint” (De Gennaro et al., 2005) because of the trait-like nature of spindles(Silverstein & Michael Levy, 1976). Previous studies have revealed that interindividual differences in spindle characteristics are related to the capacity for reasoning (i.e., the ability to identify complex patterns and relationships, the use of logic, existing knowledge, skills, and experience to solve novel problems (Fogel & Smith, 2007; Fogel & Smith, 2006; Nader & Smith, 2001, 2003). Moreover, the relationship between spindles and cognitive abilities is specific to the capacity for Reasoning, over-and-above Verbal abilities and short-term memory(Fang et al., 2017; Fogel et al., 2007). These studies have provided insight into the electrophysiological correlates of Reasoning abilities, insofar as to suggest that efficient functioning of the neural substrates that support spindle generation (e.g., thalamocortical circuitry) may be related to the capacity for these cognitive skills. Interestingly, spindle production is reduced with age (Carrier et al., 2001; Fogel et al., 2014; Fogel et al., 2017), and abnormal in developmental disorders, such as Autism (Limoges et al., 2005), learning disabilities (Shibagaki et al., 1982) and in schizophrenia (Wamsley et al., 2012). Thus, a better understanding of the neural basis of the relationship between spindles and cognitive abilities may ultimately help to better understand the significance to a variety of normal and abnormal cognitive functioning in healthy individuals and in neurological conditions. This may eventually lead to novel interventions to precisely target cases where spindle production is abnormal or non-optimal. However, it is necessary to first understand the physiological correlates of the relationship between spindles and Reasoning abilities in healthy individuals, which is the principle aim of the current study.

The association between sleep spindles and individual differences in cognitive abilities has been well documented. For example, Nader and Smith (Fogel & Smith, 2006; Nader & Smith, 2001, 2003) found that both the number of sleep spindles and sigma power (12–14 Hz) uniquely correlated with Performance IQ scores, over-and-above Verbal IQ (Fogel et al., 2007). Consistently, Bodizs and colleagues (Bódizs et al., 2005) found that spindle density was correlated with Reasoning abilities (i.e., “fluid intelligence”) measured by the Raven’s Progressive Matrices (Raven, Court, and Raven 1976). Similar studies identified a positive correlation between right-parietal fast spindles and visuospatial abilities assessed by the Rey– Osterrieth Complex Figure test (Bódizs et al., 2008), and a positive correlation between spindles and the intellectual abilities measured by the Cattell Culture Fair Intelligence Test, specifically in woman but not in men (Ujma et al., 2014). Although, a relationship in men was subsequently identified by the same group in daytime sleep (Ujma et al., 2015). Most recently, Fang and colleagues (Fang et al., 2017) used the Cambridge Brain Sciences (**CBS**) test battery (Hampshire et al., 2012) to explore if the relationship between sleep spindles and intellectual ability was a direct relationship, or whether this could be partially (or fully explained) by other spindle-related factors such as sleep quality or circadian chronotype. They found that, indeed, the relationship between spindles and Reasoning abilities was independent of sleep quality and circadian chronotype. Taken together, these studies support the notion that sleep spindles are an electrophysiological marker of cognitive abilities, and specifically, the ability to solve problems using logic and reasoning. However, the brain regions supporting the relationship between the sleep spindles and cognitive abilities are still unknown.

Only a small number of studies have employed simultaneous electroencephalography and functional magnetic resonance imaging (**EEG-fMRI**) to explore brain activations time-locked to spindles (Andrade et al., 2011; Caporro et al., 2012; Laufs et al., 2007; Schabus et al., 2007; Tyvaert et al., 2008). Spindle-related activations have been consistently found in the thalamus and the temporal lobe, for both fast spindles and slow spindles (Andrade et al., 2011; Caporro et al., 2012; Laufs et al., 2007; Schabus et al., 2007; Tyvaert et al., 2008), and activation of the cingulate cortex and motor areas have been reported to be associated with sleep spindles during NREM sleep (Andrade et al., 2011; Caporro et al., 2012). Interestingly, activation of the putamen has also been found to be correlated with spindle events (Caporro et al., 2012; Tyvaert et al., 2008) and Andrade et al. (2011) found a strong interaction between sleep spindle occurrence and hippocampal formation functional connectivity. In addition, by directly comparing fast spindles vs. slow spindles, Schabus et al. (2007) observed that slow spindles increase activations in the superior temporal gyrus while fast spindles recruit activations in the sensorimotor area, mesial frontal cortex, hippocampus, and cerebellum. Not surprisingly, given the methodological complexities and limitations of EEG-fMRI recordings during sleep, most of these studies used relatively small sample sizes (n < 15), suggesting that additional studies investigating the neural correlates of sleep spindles in a larger sample is warranted. Nonetheless, taken together, the extant literature intriguingly suggest that brain activations associated with the action of sleep spindles involve well-known spindle-generating regions (e.g., thalamic and cortical regions), as well as regions which subserve cognitive functioning and memory (e.g., hippocampal, striatal, prefrontal, motor cortical and cerebellar regions).

Interestingly, some of the regions activated during spindle events, are thought to support human cognitive abilities. For example, the thalamocortical circuitry, one of the most important neural substrates related to spindle generation (Steriade, Contreras et al.,1993; Steriade, McCormick, & Sejnowski, 1993), has been observed to be involved in reasoning abilities assessed by Raven’s Progressive Matrices test (Gray et al., 2003), the Wechsler Adult Scale of Intelligence (WAIS-III) (Wecshler, 1997), and other reasoning ability-related tasks, especially with regard to the prefrontal cortex and the thalamus (Bugg et al., 2006; Kroger, 2002; Melrose, Poulin, & Stern, 2007; Waltz et al., 1999). In addition, the basal ganglia region, especially the striatal areas (i.e. caudate and putamen), which are recruited during spindle events (Caporro et al., 2012; Tyvaert et al., 2008), have also been found to be related to cognitive functions, including reward-based learning (O’Doherty, 2004), planning (Elsinger et al., 2006), motor execution (Monchi et al., 2006), and reasoning (Melrose et al., 2007; Rodriguez-Moreno & Hirsch, 2009). Recently, Hampshire et al. (2012) employed the Cambridge Brain Sciences cognitive test battery to identify and distinguish the brain networks that support distinct cognitive abilities (e.g., Reasoning, Verbal, and Short Term Memory). It was found that the inferior frontal sulcus, the inferior parietal cortex, and the dorsal portion of the anterior cingulate / supplementary motor area activations related to Reasoning abilities and were disassociated from brain regions that related to Verbal abilities and Short Term Memory. While it is intriguing that a subset of regions which support Reasoning abilities are also regions activated with the occurrence of sleep spindles, it remains to be investigated whether spindle-related activations in these areas are correlated with interindividual differences in Reasoning abilities.

Thus, it is clear that spindle characteristics are linked to Reasoning abilities, however, the neural correlates of this relationship remain unknown. Therefore, here, using a large sample of simultaneous EEG-fMRI recordings during sleep, we sought to identify, for the first time, the neuroanatomical function correlates of the relationship between sleep spindles and Reasoning abilities. We hypothesized that the neural activation patterns, time-locked to spindles would be related to distinct cognitive abilities whereby consistent with previous cognitive and electrophysiological studies, spindle-related brain activations would be correlated with Reasoning but not STM or Verbal abilities. This will provide insight into the neural basis of the functional correlates of sleep spindles.

## RESULTS

### Cognitive abilities: Cambridge Brain Sciences Trials

Based on previous literature (Hampshire et al., 2012), the raw scores from each of the 12 subtests were Z-score normalized using the mean and standard deviation of each subtest from a large population (N = 44,600). Each test item was then weighted according to the factor loadings from Hampshire et al. (Hampshire et al., 2012) and then the respective sub-tests were averaged to create the Reasoning, STM and Verbal sub-scores and transformed to a mean of 100 and a SD of 15, so that test scores were readily comparable to results from similar studies that employed test batteries tapping into Reasoning and Verbal abilities, such as the Multidimensional Aptitude Battery - II (Fogel et al., 2007; Fogel & Smith, 2006) and other commonly used batteries of cognitive abilities (e.g., Wechsler Adult Intelligence Scale). The descriptive statistics of each subtest are shown in **Table 1.**

**Table 1.** Descriptive statistics of the 3 CBS Trials subscales (Reasoning, Short-term memory (STM) and Verbal abilities).

| **IQ Measures** | **Range** | **Mean ± SD** | **Median** |
| --- | --- | --- | --- |
| **Reasoning** | 78.84 - 108.17 | 95.65 ± 7.20 | 96.46 |
| **STM** | 84.38 - 115.33 | 101.60 ± 6.77 | 102.30 |
| **Verbal** | 88.51 - 110.92 | 99.62 ± 5.12 | 99.52 |

### Sleep Architecture

Participants slept, on average, a total of 44.20 (SD=23.84) minutes in the scanner during the experimental sleep session (**Table 2**). All N=29 participants experienced NREM2 sleep, N=20 had SWS sleep, and N=8 had rapid eye movement (REM) sleep. Given the focus of the current investigation, we did not analyze REM data. Participants had on average, a total of 334.74 (SD=212.29) total bandwidth sleep spindles at Cz during NREM sleep. Spindle parameters for all spindles at Cz (11-16 Hz), slow spindles at Fz (11-13.5 Hz) and fast spindles at Pz (13.5-16 Hz) during NREM sleep are shown in the **Table 2**.

**Table 2.** Sleep architecture and sleep spindle parameters for spindles at Fz, Cz and Pz during NREM sleep from EEG-fMRI recording sessions.

|  | **M** | **SD** |
| --- | --- | --- |
| **Sleep Architecture** |  |  |
| **Wake (min)** | 26.87 | 20.25 |
| **NREM1 (min)** | 5.84 | 4.38 |
| **NREM2 (min)** | 23.87 | 14.50 |
| **SWS (min)** | 14.77 | 17.17 |
| **NREM (min)** | 39.29 | 19.33 |
| **REM** | 17.80 | 10.76 |
| **Total Sleep** | 44.20 | 23.84 |
| **Total bandwidth (11-16 Hz) spindles at Cz** |  |  |
| **Number** | 334.74 | 212.29 |
| **Duration (sec)** | 0.49 | 0.05 |
| **Amplitude (μV)** | 27.21 | 6.43 |
| **Density** | 8.22 | 2.34 |
| **Slow spindles (11-13.5 Hz) at Fz** |  |  |
| **Number** | 249.41 | 179.40 |
| **Duration (sec)** | 0.38 | 0.06 |
| **Amplitude (μV)** | 42.71 | 9.44 |
| **Density** | 6.00 | 2.20 |
| **Fast (13.5-16 Hz) spindles at Pz** |  |  |
| **Number** | 92.48 | 69.66 |
| **Duration (sec)** | 0.38 | 0.08 |
| **Amplitude (μV)** | 21.53 | 6.01 |
| **Density** | 2.50 | 1.80 |

*Abbreviations: non-rapid eye movement sleep (NREM); stage 1 sleep (NREM1); stage 2 sleep (NREM2); slow wave sleep (SWS); rapid eye movement sleep.*

### Relationship between sleep spindles and cognitive abilities

Standard multiple linear regression revealed that, taken together, Reasoning, Short Term Memory and Verbal abilities significantly accounted for variability in spindle amplitude (F(3, 25) = 4.884, r^2^ = 0.370, p = 0.008), but not duration (F(3, 25) = 0.531, r^2^ = 0.060, p = 0.665) or density (F(3, 25) = 2.522, r^2^ = 0.232 p = 0.081) at Cz during NREM sleep (**Table 3**). Similar to previous studies (Fang et al., 2017; Fogel et al., 2007), inspection of the semipartial coefficients revealed that Reasoning ability (t(25) = 2.191, r = 0.401, p = 0.038; **Figure 1**) uniquely accounted for variability in spindle amplitude over and above STM (t(25) = 0.314, r = 0.063, p = 0.756) and Verbal (t(25) = 0.972, r = 0.191, p = 0.341) abilities. The same regression analyses were conducted for slow spindles at Fz and fast spindles at Pz, however, we did not observe any significant relationship between spindles and cognitive abilities.

**Table 3.** Multiple regression analyses of the relationship between Cambridge Brain Sciences Trials and 11-16Hz spindles at Cz during NREM sleep. See **Figure 1**.

| Overall regression effect |  |  |  |
| --- | --- | --- | --- |
| Sleep Spindle parameter | $r^2$ | $F(3,25)$ | $p$ |
| Amplitude | 0.37 | 4.884 | 0.008* |
| Duration | 0.060 | 0.531 | 0.665 |
| Density | 0.232 | 2.522 | 0.081 |
| Post-hoc effects analyses |  |  |  |
| CBS measures | Semipartial $r$ | $t(25)$ | $p$ |
| Reasoning | 0.401 | 2.191 | 0.038* |
| Verbal | 0.191 | 0.972 | 0.341 |
| STM | 0.063 | 0.314 | 0.756 |

**Figure 1.**
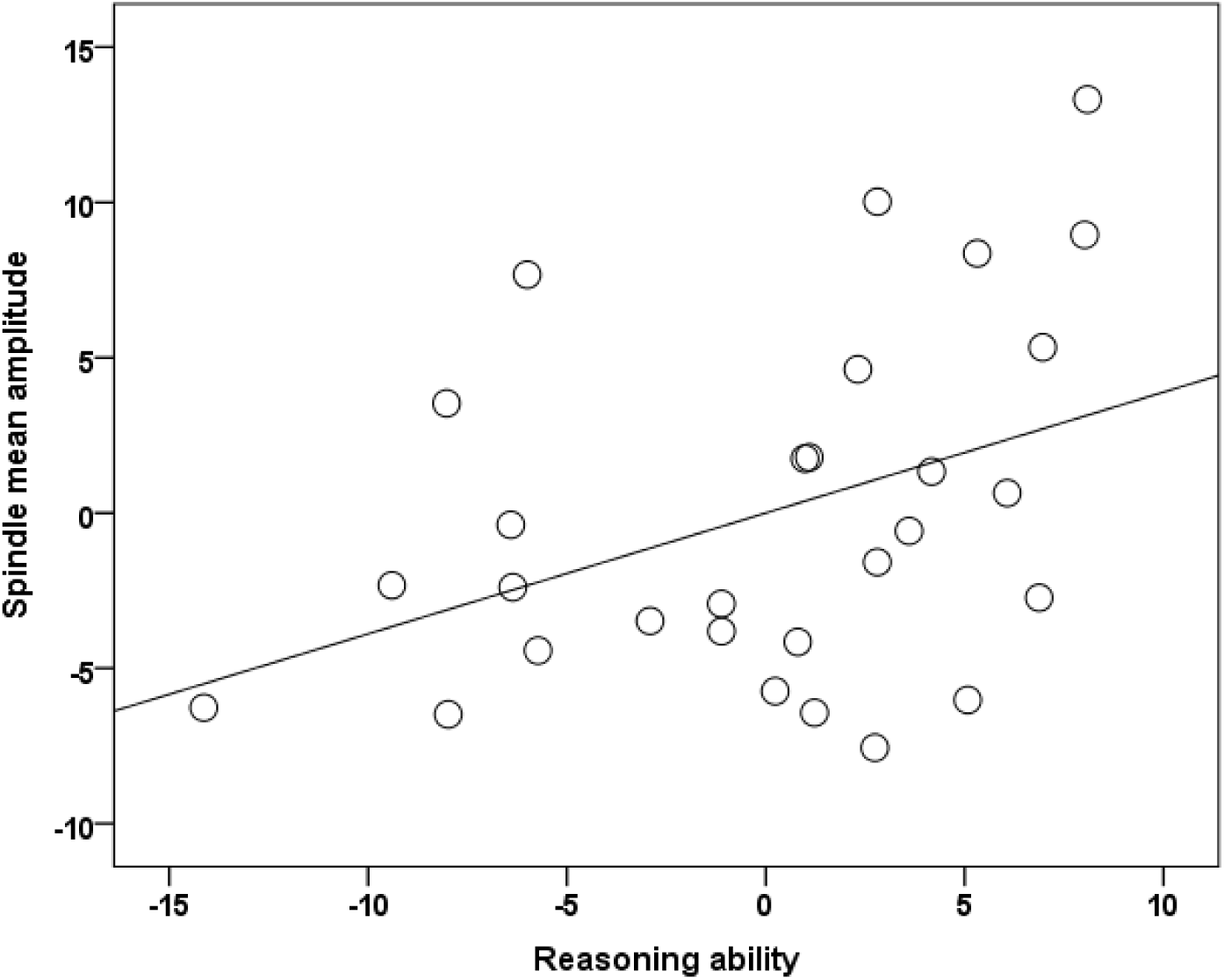
The unique relationship (i.e., semipartial correlation, r (29) = 0.401, p = 0.038) between Reasoning ability, over-and-above STM and Verbal abilities with spindle amplitude during NREM2.

### Activation of brain regions time-locked to spindles during NREM sleep

As shown in **Figure 2A**, activations time-locked to all spindles (11-16 Hz) at Cz during NREM sleep, were observed in the thalamus/midbrain, the bilateral striatum (putamen/globus pallidum and caudate), the medial frontal cortex, cerebellum, and the brain stem (cluster-level FWE corrected p < 0.05, **Table 4**). These results were statistically robust, as it is worth noting that even when a conservative whole-brain voxel-wise FWE statistical threshold correction (p < 0.05) was used, activations remained statistically significant in the thalamus/midbrain, the brainstem, the cerebellum, and the right putamen.

**Figure 2.**
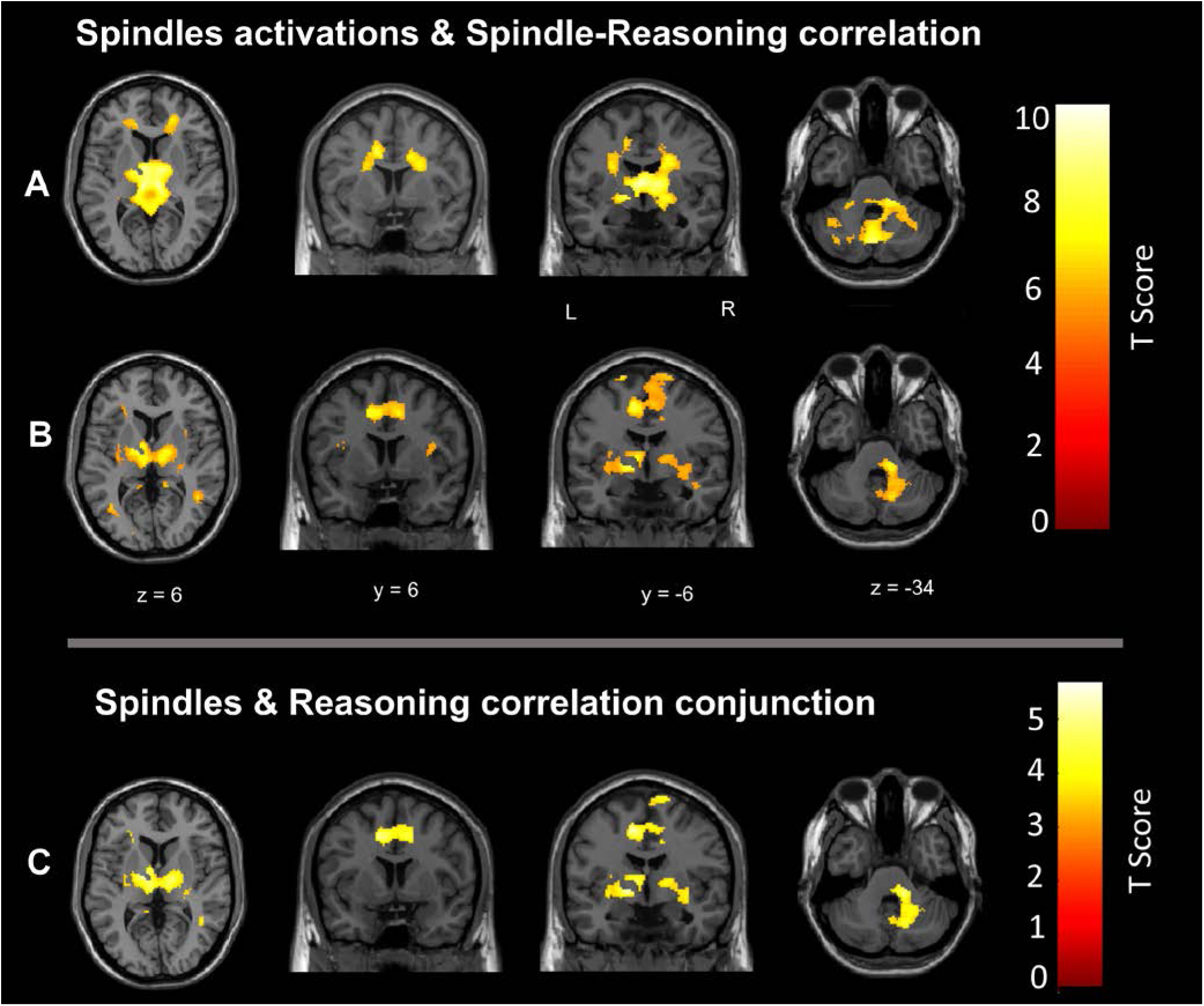
Cerebral activation time-locked to sleep spindles and correlation between spindle-related activation and Reasoning abilities. **A.** Activations time-locked to sleep spindles per se during NREM sleep. **B.** Spatial correlation maps between activations time-locked to sleep spindles and Reasoning abilities. **C.** Conjunction between A and B.

Given the two physiologically distinct spindles types (fast and slow), we also explored the brain activations time-locked to fast (13.5-16 Hz) spindles and slow (11-13.5 Hz) spindles during NREM sleep. As shown in **Figure S1**, activations time-locked to fast spindles at Pz (**Figure S1A**) and slow spindles at Fz (**Figure S1B**) were very similar in most brain regions, including the thalamus, the precuneus, and the cerebellum. There were no significant differences between fast spindle and slow spindle-related activations.

**Table 4.**
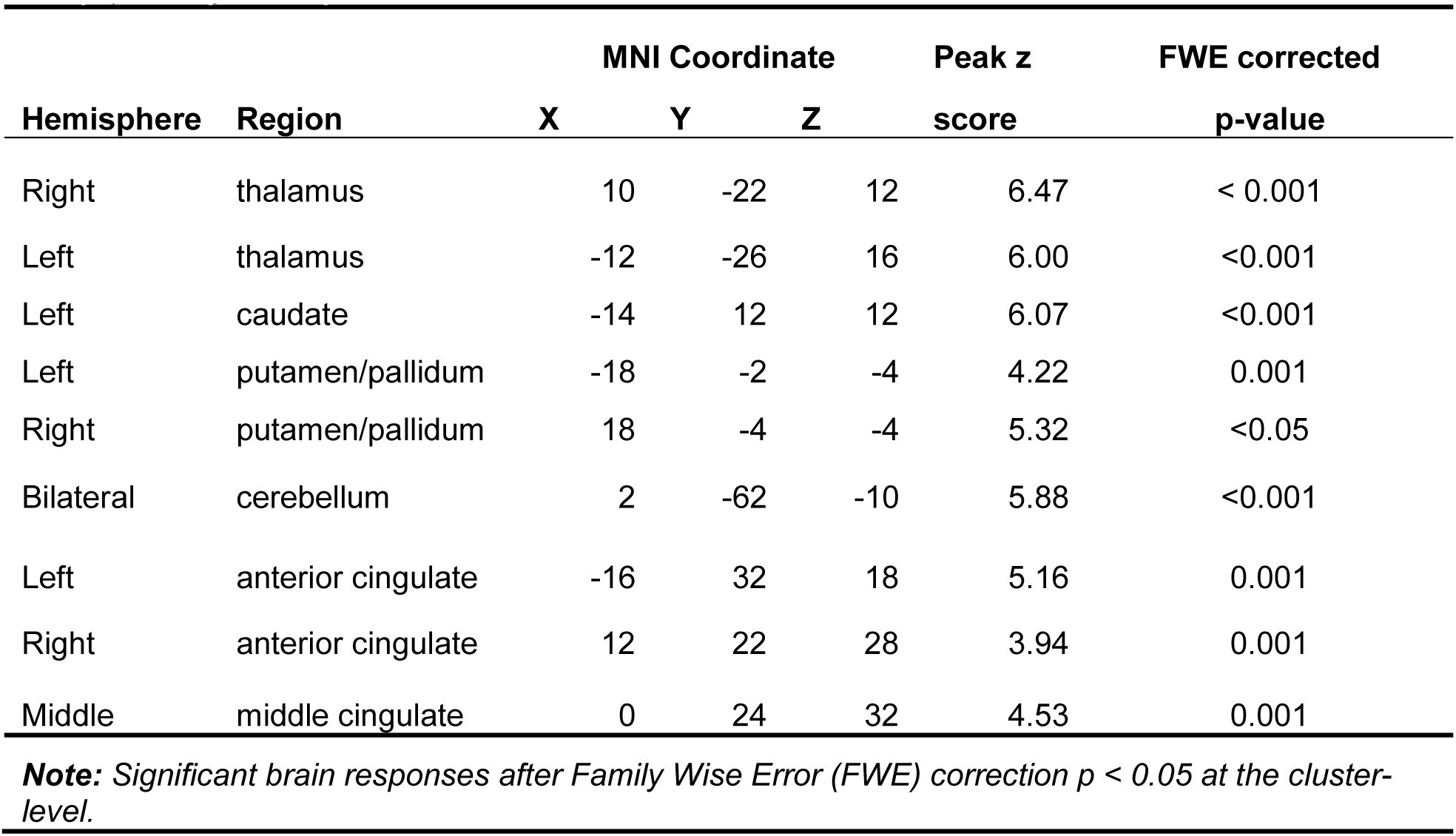
Statistically significant activations time-locked to 11-16Hz sleep spindles at Cz during NREM sleep (see **Figure 2A**).

| Hemisphere | Region | MNI Coordinate |  |  | Peak z | FWE corrected |
| --- | --- | --- | --- | --- | --- | --- |
|  |  | X | Y | Z | score | p-value |
| Right | thalamus | 10 | -22 | 12 | 6.47 | < 0.001 |
| Left | thalamus | -12 | -26 | 16 | 6.00 | <0.001 |
| Left | caudate | -14 | 12 | 12 | 6.07 | <0.001 |
| Left | putamen/pallidum | -18 | -2 | -4 | 4.22 | 0.001 |
| Right | putamen/pallidum | 18 | -4 | -4 | 5.32 | <0.05 |
| Bilateral | cerebellum | 2 | -62 | -10 | 5.88 | <0.001 |
| Left | anterior cingulate | -16 | 32 | 18 | 5.16 | 0.001 |
| Right | anterior cingulate | 12 | 22 | 28 | 3.94 | 0.001 |
| Middle | middle cingulate | 0 | 24 | 32 | 4.53 | 0.001 |

**Note:** Significant brain responses after Family Wise Error (FWE) correction $p < 0.05$ at the cluster-level.

### Correlation between cognitive abilities and brain activations time-locked to spindles

To examine the neural correlates of the relationship between sleep spindles and cognitive abilities, we conducted whole-brain spatial correlation analyses between brain activation maps time-locked to all spindles at Cz and the scores on the three cognitive factors (Reasoning, STM, and Verbal abilities) assessed by the Cambridge Brain Sciences tests. As shown in **Figure 2B**, Reasoning ability was significantly correlated with spindle-related activation maps in the thalamus, bilateral putamen, brainstem/pons anterior cingulate cortex, the middle cingulate cortex, the paracentral lobe, the posterior cingulate cortex, the precuneus, and bilateral temporal lobe (see **Table 5**).

**Table 5.**
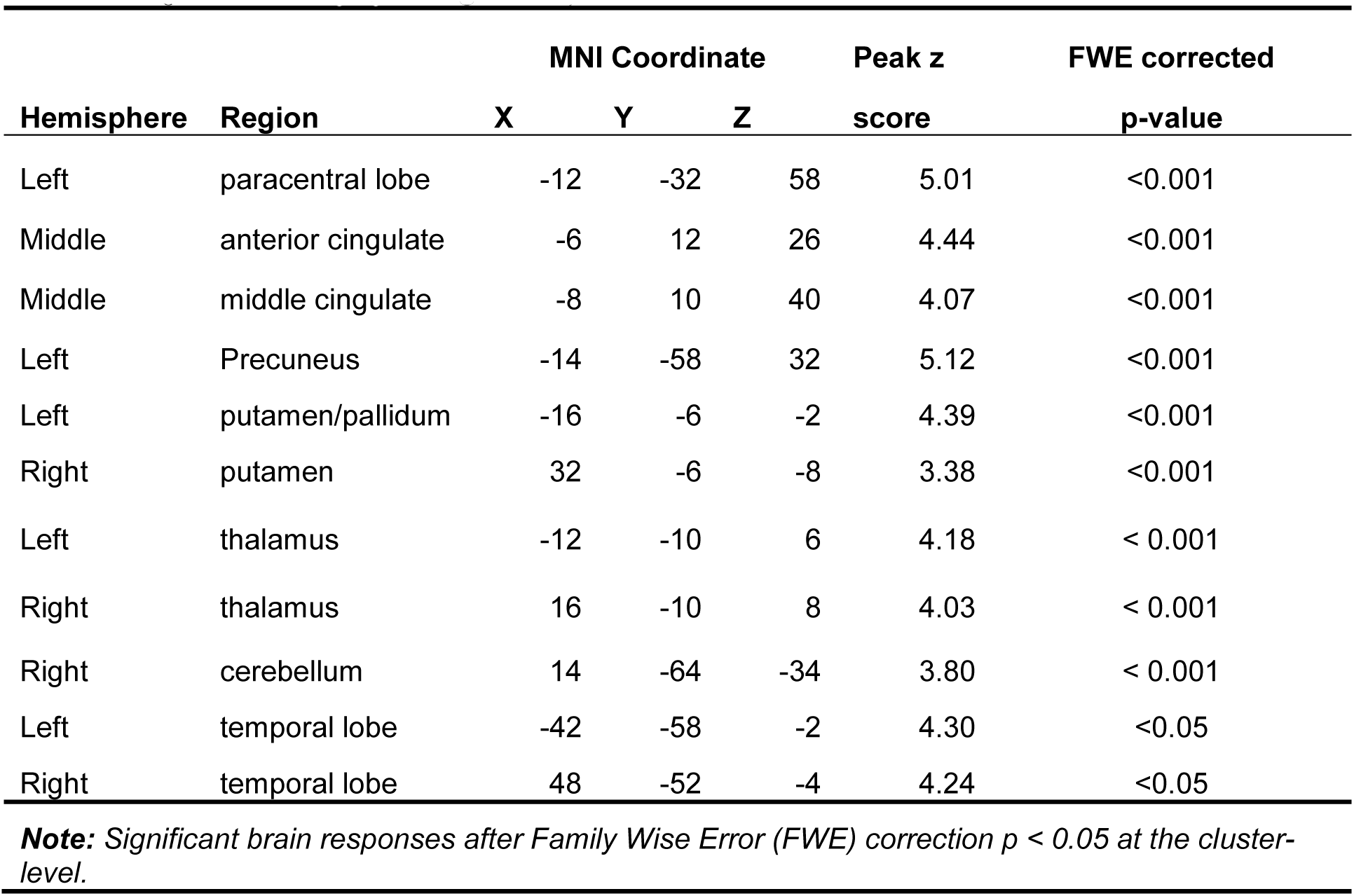
Whole brain correlations between Reasoning ability and 11-16Hz spindle-related activations at Cz during NREM sleep. (see **Figure 2B**)

| Hemisphere | Region | MNI Coordinate |  |  | Peak z | FWE corrected |
| --- | --- | --- | --- | --- | --- | --- |
|  |  | X | Y | Z | score | p-value |
| Left | paracentral lobe | -12 | -32 | 58 | 5.01 | <0.001 |
| Middle | anterior cingulate | -6 | 12 | 26 | 4.44 | <0.001 |
| Middle | middle cingulate | -8 | 10 | 40 | 4.07 | <0.001 |
| Left | Precuneus | -14 | -58 | 32 | 5.12 | <0.001 |
| Left | putamen/pallidum | -16 | -6 | -2 | 4.39 | <0.001 |
| Right | putamen | 32 | -6 | -8 | 3.38 | <0.001 |
| Left | thalamus | -12 | -10 | 6 | 4.18 | < 0.001 |
| Right | thalamus | 16 | -10 | 8 | 4.03 | < 0.001 |
| Right | cerebellum | 14 | -64 | -34 | 3.80 | < 0.001 |
| Left | temporal lobe | -42 | -58 | -2 | 4.30 | <0.05 |
| Right | temporal lobe | 48 | -52 | -4 | 4.24 | <0.05 |

**Note:** Significant brain responses after Family Wise Error (FWE) correction $p < 0.05$ at the cluster-level.

Since Reasoning ability was highly inter-correlated with Verbal ability (r = 0.596. p = 0.001), and marginally correlated with STM ability (r = 0.357, p = 0.058), to ensure that the relationship between Reasoning ability and the spindle-related activations was not accounted for by Verbal or STM abilities, we also examined the correlations between spindle-related activations and Short Term Memory, and also Verbal abilities, respectively. However, no significantly correlated activations were observed between Short Term Memory or Verbal abilities and the spindle-related activations. This demonstrates that a subset of spindle-related activations was specifically related to Reasoning abilities, but not to Short Term Memory or Verbal abilities. This is consistent with, and provides physiological support for the current and previous studies (Bódizs et al., 2005; Fang et al., 2017; Fogel et al., 2007; Schabus et al., 2006; Ujma et al., 2014, 2015), demonstrating that Reasoning abilities are uniquely correlated to sleep spindles. The same whole-brain spatial correlation analyses were conducted for fast spindle and slow spindle activation maps, however, we did not observe significant correlations between any cognitive ability and the activation maps for each individual spindle type.

From **Figure 2**, we can see that there were several overlapping regions between the spindle activation maps (**Figure 2A**) and the maps that show activations time-locked to spindles that were correlated with Reasoning abilities (**Figure 2B**). The conjunction (at p < 0.001 using the conjunction null) between the spindle activation maps and the Reasoning-related spindle correlation maps (**Figure 2C)**, show several regions were consistently high and jointly activated in both the spindle and Reasoning-spindle correlation maps, including the thalamus, medial frontal cortex, bilateral putamen, and the cerebellum (**Table 6**).

**Table 6.**
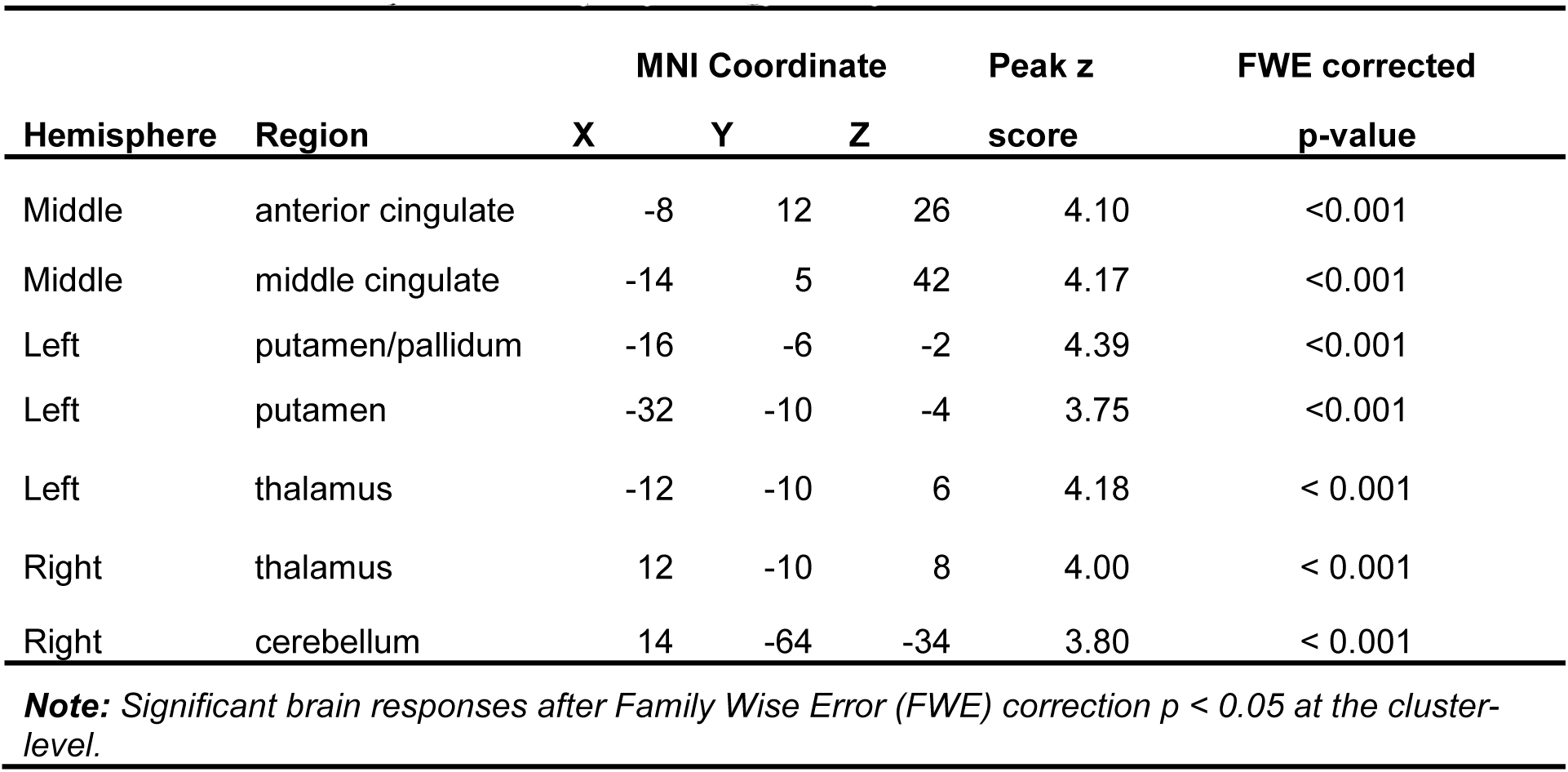
Conjunction between the spindle-related activation maps and the spindle-related activations correlated with reasoning abilities maps. (see **Figure 2C**)

Finally, to ensure that activations time-locked to spindles were specific to spindles per se, and not to some general epiphenomena of NREM sleep, a separate analysis investigated activations time-locked to the same number of randomly distributed onsets during NREM sleep, instead of onsets aligned to spindles events. The results revealed only a small single cluster at the left frontal lobe (peak coordinate: -28, -2, 68; uncorrected p < 0.005), which did not overlap with the activations time-locked to spindles (**Figure S2**), and did not survive correction for multiple comparisons, suggesting that this activation is non-specific to NREM sleep, and likely spurious. No correlation was observed between Reasoning ability and the uncorrected random onsets map. This suggests that the reactivations reported here, are specifically related to spindle events, and not simply to NREM sleep in general.

## DISCUSSION

Sleep supports normal human cognitive performance, such as attention, language, reasoning, decision making, learning and memory (for review, see Alhola & Polo-Kantola 2007; Diekelmann 2014; Diekelmann & Born 2010; Goel et al. 2009; Harrison & Horne 2000). Previous EEG studies have identified sleep spindles as a biological marker of cognitive abilities, and in particular, reasoning abilities (Bódizs et al., 2005; Fang et al., 2017; Fogel et al., 2007; Fogel & Smith, 2011; Schabus et al., 2006; Ujma et al., 2014, 2015). Only a few EEG-fMRI studies have explored the brain activations correlated with sleep spindles (Andrade et al., 2011; Caporro et al., 2012; Laufs et al., 2007; Schabus et al., 2007; Tyvaert et al., 2008). Interestingly, some of these regions are also known to support reasoning abilities. However, the neuroanatomical functional correlates of the relationship between spindles and Reasoning abilities are unknown. Here, we identified the neural activation patterns time-locked to spindles that are correlated to cognitive abilities. Using a large sample of simultaneous EEG-fMRI sleep recordings, the results of the present study support three main findings: (1) similar to previous studies (Fang et al., 2017; Fogel et al., 2007), spindles detected at Cz (11-16Hz) during NREM sleep were related to Reasoning but not Short Term Memory or Verbal abilities, (2) similar to previous studies (Andrade et al., 2011; Caporro et al., 2012; Laufs et al., 2007; Schabus et al., 2007; Tyvaert et al., 2008), activations time-locked to spindles were observed in the thalamus, bilateral striatum, middle cingulate cortex, and cerebellum, and (3) Reasoning abilities were correlated with spindle-related activations in a subset of these regions including the thalamus, bilateral striatum, medial frontal gyrus, middle cingulate cortex, and precuneus. These results are specific to spindles *per se*, and cannot be attributed to some epiphenomena during NREM sleep; given that these results were not observed when random onsets during NREM sleep were used instead of onsets time-locked to spindle events. Altogether, our results identified for the first time, the neural correlates of the relationship between spindles and Reasoning abilities.

### Spindle-related activation of thalamocortical circuitry

Consistent with previous EEG-fMRI studies of spindle-related activations (Andrade et al., 2011; Caporro et al., 2012; Laufs et al., 2007; Schabus et al., 2007; Tyvaert et al., 2008), our results identified and confirmed the brain regions associated with spindle events during NREM sleep in both cortical (including the media prefrontal, anterior cingulate cortex, and middle cingulate cortex), and subcortical areas (including the thalamus and bilateral caudate, putamen, and pallidum), indicating the role of cortico-thalamic-basal ganglia circuitry in spindle generation. In addition, positron emission tomography (PET) studies have shown changes in regional cerebral blood flow in the thalamus related to sleep spindles (Hofle et al., 1997). These human neuroimaging findings are supported by a large body of animal studies, which at the cellular level, suggest that spindles reflect oscillatory activity in widespread thalamocortical circuits, and involve complex interactions between reticular, thalamocortical and pyramidal cells (Steriade, 2005). Classically, spindle generation was shown to be maintained by synchronized firing in the reticular-thalamocortical-reticular circuit (Steriade, Nunez, & Amzica, 1993; von Krosigk et al., 1993). More recent evidence (Bonjean et al., 2011) however, suggests that corticothalamic input initiates spindles by triggering spike bursts in the reticular nucleus and are terminated by desynchronization of thalamic and cortical neuronal firing. Thus, taken together, animal and recent human neuroimaging studies, including the current study, supports the involvement of thalamocortical circuitry in spindle generation.

The results of the current study identified a correlation between Reasoning, but not Short Term Memory or Verbal abilities with spindle-related brain activations in thalamocortical circuitry, especially the thalamus and the prefrontal cortex (PFC) region, which are thought to be implicated in modulation of cognitive performance (Blair, 2006; Ferguson & Gao, 2015; Mitchell & Chakraborty, 2013). Spindles and the thalamus have been shown to be related to human intellectual abilities (Fangmeier et al., 2006; Melrose et al., 2007). The thalamus, especially the mediodorsal thalamus has been reported to be related to the fluid intelligence (Van der Werf et al., 2003; Van Der Werf et al., 2000) particularly for the extrapolation component process of inductive reasoning (Jia et al., 2011; Liang et al., 2014) and other higher-level cognition (e.g., problem solving, working memory, goal direct seeking) (Karatekin et al., 2000; Mitchell & Chakraborty, 2013; Schiff et al., 2002; Shirvalkar et al., 2006). Lesions studies in both humans (De Witte et al., 2011; Kubat-Silman et al., 2002; Little et al., 2010) and non-human primates (for review, see Mitchell & Chakraborty 2013) have revealed that thalamic damage impairs various broadly defined aspects of cognitive performance, including discrimination, memory, learning, attention and other neuropsychological behaviors. Other neuroanatomical studies (Bohlken et al., 2014) found that only thalamic volume was significantly correlated with general intellectual functioning. In addition, at least one study identified structural and functional abnormalities in the thalamus in adults with reduced intellectual functioning who experienced prenatal exposure to alcohol (Clark et al., 2000). Thus, suggesting that integrity and functioning of the neural circuitry involved in spindle generation support intellectual abilities, and in particular Reasoning abilities.

The prefrontal cortex has been defined as the projection area of the mediodorsal thalamus (Behrens et al., 2003), and the prefrontal-thalamic loop plays a critical role in various higher-order cognitive processes, especially executive function (for reviews see Baxter 2013; Ferguson & Gao 2015; Funahashi 2013; Mitchell & Chakraborty 2013; Watanabe & Funahashi 2012). A large body of literature has identified the role of the prefrontal cortex area in fluid intelligence and Reasoning (Coricelli & Nagel, 2009; Duncan, 2000; Gray et al., 2003; Melrose et al., 2007; Sandman et al., 2014; Waltz et al., 1999). For example, patients with damage to the prefrontal cortex exhibited a selective and catastrophic deficit for both deductive and inductive reasoning tasks (Waltz et al., 1999). In addition, Gray et al., (2003) found that individuals with higher fluid intelligence have greater activations in the prefrontal cortex. Coricelli and Nagel (2009) have shown that reasoning abilities correlate with neural activity in the medial prefrontal cortex. Additionally, at least one neuroanatomical MRI study employing voxel-based morphometry has revealed a positive correlation between gray matter intensity in the medial prefrontal cortex and reasoning abilities assessed by Cattell’s Culture Fair Intelligence Test, and also the WAIS-R (Gong et al., 2005). Taken together, these findings suggest that the thalamus and prefrontal cortex region supports Reasoning abilities.

### Spindle-related activation of the basal ganglia

Consistent with previous results (Caporro et al., 2012; Tyvaert et al., 2008), the present study shows that the basal ganglia, including striatal areas (caudate and putamen) and the globus pallidus were recruited during spindle events. The basal ganglia are primarily known for playing a role in cognitive functions (for reviews see Burgaleta et al. 2014; Chakravarthy et al. 2010; Leisman et al. 2014; Doyon et al. 2009) such as action selection, reward-based learning, motor sequence learning, planning (Elsinger et al., 2006) and motor execution (Monchi et al., 2006). In addition, several studies have observed robust activations in the basal ganglia for reasoning-related tasks compared to other cognitive tasks, including the caudate nucleus, putamen and globus pallidus (Ferguson & Gao, 2015; Melrose et al., 2007; Rodriguez-Moreno & Hirsch, 2009). Taken together, these findings complement the results of the current study whereby activation of the putamen time-locked spindles was correlated with Reasoning abilities. The Reasoning subtests of the Cambridge Brain Sciences Trials, consists of tasks requiring planning (Shallice, 1982), spatial rotation (Silverman et al., 2000), and visuomotor ability (Folstein et al., 1975). Sandman et al., (2014) has reported that the morphometry of the putamen was associated with performance on reasoning-related subtests of the WAIS including block design, matrix reasoning and perceptual index in preadolescent children. These findings suggest that the function and structure of the basal ganglia are related to Reasoning abilities. The current study suggests that interindividual differences in spindle-related activation of these regions are related to Reasoning ability.

### Spindle-related activation of the cerebellum

Similar to previous studies (Schabus et al., 2007), here, we also observed spindle-related activation of the cerebellum which was also correlated with Reasoning abilities. Many studies overlook cognition-related activity in the cerebellum, although this area is responsible for modulating thalamic activity through direct cerebello-thalamic projections, which may be related to spindle generation (Calzavara et al., 2005; Shouse & Sterman, 1979). In addition, it has been suggested that the cerebellum supports cognitive functions (for reviews, see Gordon 2007; Rapoport et al. 2000; Stoodley 2012) such as response reassignment during a complex task (Bischoff-Grethe et al., 2002), decision making (Blackwood et al., 2004), associative learning (Logan & Grafton, 1995), adaptation (Krakauer & Mazzoni, 2011), and executive function (Bellebaum & Daum, 2007; Tomasi et al., 2007). Importantly, the cerebellum is activated during the deductive reasoning processing (Goel et al., 2000; Goel & Dolan, 2004) supporting our finding that reasoning-related functions supported by the cerebellum is reflected during sleep, time-locked to spindles.

Unlike previous studies, we did not observe spindle-related activation of the medial temporal lobe (Andrade et al., 2011; Caporro et al., 2012; Laufs et al., 2007; Schabus et al., 2007; Tyvaert et al., 2008). In addition, Schabus et al., (2007) and Andrade et al. (2011) reported brain activation differences between fast spindles and slow spindles, particularly in hippocampal regions. However, here, we observed similar activation patterns for both spindle types and no significant differences between spindle types. However, we did find medial frontal activation for slow, but not fast spindle events. Despite having a large proportion of participants who slept for an adequate amount of NREM sleep, there may not have been an adequate number of each spindle type when categorized orthogonally for sufficient power, or perhaps not enough instersubject variability to detect any relationship to cognitive abilities. Moreover, sleep was recorded only from the first couple of hours of the night, where the relationship between Reasoning abilities and spindles has been found to be much less robust than the later part of the night (Fogel et al., 2007). This might also help explain that when separated into fast spindles and slow spindles, we did not observe significant relationship with Reasoning abilities in each spindle subtype. Lastly, due to the limited duration, and high intersubject variability of sleep in the scanner, we did not have sufficient SWS to test whether a different pattern of results was observed during SWS.

The clinical significance and applications of the relationship between spindles and cognitive abilities is yet to be realized. Deficient or dysfunctional spindle generation may be associated with compromised intellectual functioning. More specifically, it has been suggested that deficient gating mechanisms of thalamocortical circuitry (Bixler,1968) may explain abnormal spindle production in children with mental disability (Gibbs & Gibbs, 1962; Shibagaki & Kiyono, 1983). Moreover, the present study is an important first step which may lead to the development of novel interventions utilizing spindle-enhancing neuromodulatory techniques (e.g., neurofeedback, transcranial direct current stimulation, pharmacological) to improve daytime cognitive performance and explore the physiological mechanisms which support the function of sleep for memory and cognitive performance. Such an approach could target cognitive deficits, in cases where spindle production is abnormal such as in learning disabilities (Gibbs & Gibbs, 1962; Shibagaki & Kiyono, 1983), below normal cognitive functioning (Fogel & Smith, 2011), normal, healthy aging (Carrier et al., 2001; Fogel et al., 2014; Fogel et al., 2017), developmental disorders (Limoges et al., 2005) and in schizophrenia (Wamsley et al., 2012).

Here, we investigated what neural substrates support cognitive strengths and weaknesses. There are considerable interindividual differences in sleep spindles, which are very trait-like(Gaillard & Blois, 1981; Silverstein & Michael Levy, 1976). While the neural circuitry and generating mechanisms of spindles are well-understood, the neurophysiological basis of the relationship between spindles and cognitive abilities remain to be fully elucidated. In summary, our results show for the first time the neuroanatomical functional correlates of the relationship between sleep spindles and intellectual abilities. In particularly, our study found that the extent of the activation of the prefrontal cortex, basal ganglia, cerebellum and the thalamus time-locked to sleep spindles was correlated with interindividual differences in Reasoning, but not Verbal or STM abilities. Thus, spindles may serve as an electrophysiological marker of brain activations in regions which support the ability to employ reasoning to solve problems and apply logic in novel situations.

## METHODS

### Participants

A total of 35 healthy right-handed adults (20 female) between 20-35 years old (M = 25.6, SD = 3.6), were recruited to participate in this study. All participants were non-shift workers and medication-free, had no history of head injury or seizures, had a normal body mass index (<25), and did not consume excessive caffeine, nicotine or alcohol. To be included, interested participants had to score <10 on the Beck Depression (Beck et al., 1974) and the Beck Anxiety (Beck et al., 1988) inventories and have no history or signs of sleep disorders indicated by the Sleep Disorders Questionnaire (Douglass et al., 1994). All participants were required to keep a regular sleep-wake cycle (bed-time between 22h00-24h00, wake-time between 07h00-09h00) and to abstain from taking daytime naps at least 7 days prior to and throughout participation in the study. Compliance with this schedule was assessed using both sleep diaries and wrist actigraphy (Actiwatch 2, Philips Respironics, Andover, MA, USA) worn on the non-dominant wrist. All participants met the MRI safety screening criteria. In addition, participants were given a letter of information, provided informed written consent before participation, and were financially compensated for their participation. This research was approved by the Western University Health Science research ethics board.

Sample sizes were determined a-priori based on previous studies, and power calculated, where possible using G*Power for Mac version 3.1 (Faul, Erdfelder, Buchner, & Lang, 2009; Faul, Erdfelder, Lang, & Buchner, 2007). Based on the most comparable simultaneous EEG-fMRI studies (Andrade et al., 2011; Caporro et al., 2012; Laufs et al., 2007; Schabus et al., 2007; Tyvaert et al., 2008), previous studies have employed sample sizes N<15. A recent study by our group using the same cognitive tests as the current study (Fang et al., 2017) found robust associations between spindles and cognitive abilities in a sample size of N=24, replicating previous findings in smaller samples (e.g., N<12: Fogel et al., 2007; Fogel & Smith, 2006). Based on power calculation for correlation with p(2-tailed) = 0.05 (b = 0.20, effect size = 0.56) from (Fang et al., 2017), an N=22 was required. Thus, N=29 subjects included in this study was considered to provide adequate statistical power for the main effects of interest.

### Experimental procedure

Each participant underwent a screening/orientation session one week prior to the experimental sleep session. All participants completed the CBS test battery online at least 3 days prior to the experimental session. The experimental sleep session took place between 21h00 and 23h00, during which time simultaneous EEG-fMRI was recorded while participants slept in the scanner. To be included in the analyses, participants were required to sleep for a period of at least 5 minutes of uninterrupted NREM sleep during the sleep session. This was considered to be the minimum amount of data necessary for EEG and fMRI data analysis purposes, and to ensure a minimum duration, quality and continuity of sleep. It should be noted that all subjects had at least 14.67 minutes of sleep, with at least 63 total bandwidth spindles (11-16 Hz) at Cz. Importantly, the average duration of NREM sleep was 39.29 minutes, with an average of 334.74 total bandwidth spindles (11-16 Hz) at Cz. Following the sleep session, participants were allowed to sleep in the nearby sleep laboratory for the remainder of the night.

Of the 35 participants who met the inclusion criteria, only 5 participants did not meet the 5-minute consolidated NREM sleep criteria for the sleep session. As well, one participant did not complete the Cambridge Brain Sciences Trials test battery. In total, 29 participants (M age = 23.97, SD = 3.83, 17 female) were included in the final data analyses.

### Cognitive ability test

The Cambridge Brain Sciences test battery (Hampshire et al., 2012) was used to assess participants’ cognitive abilities. Cambridge Brain Sciences is a web-based battery of 12 cognitive tests that assesses a broad range of cognitive abilities including reasoning, problem solving, planning, attention, and memory. A recent study, based on scores from a populationsized pool of 44,600 participants, revealed three factors that govern performance across the Cambridge Brain Sciences subtests. These factors have been described as “Reasoning”, “Short Term Memory” and “Verbal” ability (Hampshire et al., 2012). The descriptive statistics of each subtest score are shown in **Table 1.**

### Polysomnographic Recording and Analysis

#### Recording Parameters

EEG was recorded using a 64-channel magnetic resonance (MR)-compatible EEG cap which included one electrocardiogram (ECG) lead (Braincap MR, Easycap, Herrsching, Germany) and two MR-compatible 32-channel amplifiers (Brainamp MR plus, Brain Products GmbH, Gilching, Germany). EEG caps included scalp electrodes referenced to FCz. Two bipolar electrocardiogram (ECG) recordings were taken from V2-V5 and V3-V6 using an MR-compatible 16-channel bipolar amplifier (Brainamp ExG MR, Brain Products GmbH, Gilching, Germany). Using high-chloride abrasive electrode paste (Abralyt 2000 HiCL; Easycap, Herrsching, Germany), electrode-skin impedance was reduced to < 5 KOhm. To reduce movement-related EEG artifacts, participants' heads were immobilized in the MRI head-coil using foam cushions. EEG was digitized at 5000 samples per second with a 500-nV resolution. Data were analog filtered by a band-limiter low pass filter at 500 Hz and a high pass filter with a 10-sec time constant corresponding to a high pass frequency of 0.0159 Hz. Data were transferred via fiber optic cable to a personal computer where Brain Products Recorder Software, Version 1.x (Brain Products, Gilching, Germany) was synchronized to the scanner clock. EEG was monitored online with Brain Products RecView software using online artifact correction. Sleep stages were scored in accordance with standard criteria (Iber et al., 2007) using the “VisEd Marks” toolbox (https://github.com/jadesjardins/vised_marks) for eeglab (Delorme & Makeig, 2004). Automatic spindle detection was carried out using a previously published (Fogel et al., 2014; Fogel et al., 2015) and validated (Ray et al., 2015) method employing EEGlab-compatible (Delorme & Makeig, 2004) software (github.com/stuartfogel/detect_spindles) written for MATLAB R2014a (The MathWorks Inc., Natick, MA). The detailed processing steps and procedures are reported elsewhere (Ray et al., 2015) and are thus presented only briefly here. The EEG data were initially downsampled to 250 Hz. The detection was performed at Fz, Cz and Pz derivations. The spindle data were extracted from movement artifact-free, NREM stage 2 sleep epochs. The detection method (Ray et al., 2015) used a complex demodulation transformation of the EEG signal with a bandwidth of 5 Hz centered about a carrier frequency of 13.5 Hz (i.e., 11–16 Hz) (Iber et al., 2007). To utilize a fixed amplitude detection threshold, but still account for individual differences in spindles, each data point was transformed into a z-score using the mean and standard deviation derived from a 60-sec sliding window. Events (spindle onsets, peaks, and offsets) were then detected on the transformed signal with a z-score threshold of z = 2.33, corresponding to the 99^th^ percentile. The dependent variables of interest extracted from this method include spindle amplitude, spindle duration, and spindle density (number of spindles per minute of NREM sleep) for each participant and at each derivation (Fz, Cz and Pz). Spindles were categorized so that they were orthogonal (non-overlapping detections) at the scalp locations where they predominate topographically (Jobert et al., 1992; Werth et al., 1997; Zeitlhofer et al., 1997) as slow spindles (11–13.5 Hz) at Fz, total bandwidth spindles (11-16 Hz) at Cz, and fast spindles (13.5–16 Hz) at Pz (**Table 2**). Despite having no minimum detection criteria, the detection method employed here did not detect spindles lower than 0.2 sec, as found in a previous validation study (Ray et al., 2015).

### MRI Imaging Acquisition and Analysis

#### Recording Parameters

Brain images were acquired using a 3.0T TIM TRIO magnetic resonance imaging system (Siemens, Erlangen, Germany) and a 64-channel head coil. In all participants, a structural T1-weighted MRI image was acquired using a 3D MPRAGE sequence (TR = 2300 ms, TE = 2.98 ms, TI = 900 ms, FA = 9°, 176 slices, FoV = 256×256 mm^2^, matrix size = 256×256×176, voxel size = 1×1×1 mm^3^). Multislice T2*-weighted fMRI images were acquired during the sleep session with a gradient echo-planar sequence using axial slice orientation (TR = 2160 ms, TE = 30 ms, FA = 90°, 40 transverse slices, 3 mm slice thickness, 10% inter-slice gap, FoV = 220×220 mm^2^, matrix size = 64×64×40, voxel size = 3.44×3.44×3 mm^3^). Importantly, the sequence parameters were chosen so that the gradient artifact would be time stable, and the lowest harmonic of the gradient artifact (18.52 Hz) would occur outside the spindle band (11-16 Hz). This was achieved by setting the MR scan repetition time to 2160 ms, such that it matched a common multiple of the EEG sample time (0.2 ms), the product of the scanner clock precision (0.1 μs) and the number of slices (40 slices) used (Mulert & Lemieux, 2009).

### Image Preprocessing

Functional images were preprocessed and analyzed using SPM8 (http://www.fil.ion.ucl.ac.uk/spm/software/spm8/; Welcome Department of Imaging Neuroscience, London, UK) implemented in MATLAB (ver. 8.5 R2015a) for Windows (Microsoft, Inc. Redmond, WA). Functional scans of each session were realigned using rigid body transformations, iteratively optimized to minimize the residual sum of squares between the first and each subsequent image separately for each session. A mean realigned image was then created from the resulting images. The structural T1-image was coregistered to this mean functional image using a rigid body transformation optimized to maximize the normalized mutual information between the two images. Coregistration parameters were then applied to the realigned blood-oxygen-level dependent (BOLD) time series. The coregistered structural images were segmented into grey matter, white matter and cerebrospinal fluid. An average subject-based template was created using DARTEL in SPM8. All functional and anatomical images were spatially normalized using the resulting template, which was generated from the structural scans. Finally, spatial smoothing was applied on all functional images (Gaussian kernel, 8 mm full-width at half-maximum (FWHM).

### Sleep sessions

For data acquired during the simultaneous EEG-fMRI sleep recordings, within-session series of consecutive fMRI volumes sleep stage scored as NREM stage 2 sleep according to standard criteria (Iber et al., 2007) by an expert, registered polysomnographic technologist were selected from the complete fMRI time series of sleep session. To be included in the fMRI analysis, the EEG had to be visibly movement artifact-free and be a segment of uninterrupted sleep longer in duration than 55 volumes (i.e., ∼120 seconds or longer; corresponding to the minimum amount of sleep that was needed to perform the automated spindle detection), resulting in the inclusion of 36% of the total recorded data (i.e., 11,466 of 31,852 MRI volumes during NREM stage 2 sleep). Each time series corresponding to NREM stage 2 sleep that met these criteria, were entered into the general linear model (GLM) as a separate session so that no gaps existed in the design matrix. For each participant, brain responses were estimated in an event-related design using a fixed-effects GLM including responses time-locked to spindle events (11-16 Hz) detected at Cz, slow spindles (11-13.5 Hz) detected at Fz, and fast spindle events (13.5-16 Hz) detected at Pz. Consistent with similar previous studies (Andrade et al., 2011; Bergmann et al., 2011; Dang-Vu et al., 2008; Schabus et al., 2007), the vectors, including spindle events, were convolved with the canonical hemodynamic response function (HRF), as well as with its temporal and dispersion derivatives. Nuisance variables in the model included: the movement parameters estimated during realignment (translations in x, y, and z directions and rotations around x, y, and z axes), the squared value of the movement parameter, the first derivative of each movement parameter, and the square of the first derivative of each movement parameter, as well as, to the mean white matter intensity and the mean cerebral spinal fluid intensity for each participant. Slow wave activity is a defining characteristic of NREM sleep (Iber et al., 2007), but is related to spindle generation (Mölle et al., 2011; Siapas & Wilson, 1998). This activity was accounted for by including spectral power (μV^2^) in the delta band (0.5-4 Hz) for each TR window (2160 ms) as a variable of no interest, convolved with the hemodynamic response function. Slow drifts were removed from the time series using a high pass filter with a cut-off period of 128 seconds. Serial correlations in the fMRI signal were estimated using an autoregressive (order 1) plus white noise model and a restricted maximum likelihood (ReML) algorithm. These analyses generated statistical parametric t maps [(SPM(T)]. The resulting contrast images were then smoothed (FWHM 6 mm Gaussian Kernel) and entered into a second-level analysis.

The resulting group-level analysis consisted of one sample t-tests for each contrast of interest (i.e., all spindle events, fast spindle events, and slow spindle events). To investigate the relationship between the magnitude of the spindle-dependent activation and the cognitive abilities assessed by the CBS Trials, cognitive test scores for each subtest (i.e., Reasoning, Verbal, and Short Term Memory) were entered as covariates of interest in the described GLM. These activation maps constituted maps of the t-statistic [SPM(t)] testing for the main effect for each contrast of interest. Statistical inferences were performed at a threshold of p<0.05, family wise error (FWE) corrected at the cluster level.

### Overlap between spindle-related maps and reasoning-spindle correlation maps

To illustrate the overlap of activations between the spindle-related activation maps and reasoning-spindle correlation maps, the conjunction was taken as the minimum t-statistic using the conjunction hypothesis (Friston et al., 2005; Nichols et al., 2005) over: (1) a t-map testing for the main effect of the spindle events during the sleep session, and (2) a t-map testing for the main effect of the correlation between the Reasoning ability and spindle events. These two statistical maps were thresholded at p < 0.05, FWE corrected at the cluster level.

Finally, to confirm that activations time-locked to spindles and correlated with Reasoning abilities were not simply an epiphenomenon of NREM sleep, we generated the same number of random events as sleep spindles in each segment of NREM sleep for all participants during the sleep session. These random onsets did not overlap with any spindle events. We conducted the exact same GLM as for actual spindle onsets, with the only difference being that the randomly generated onsets were included in the model, as opposed to the spindle onsets.

## ACKNOWLEDGEMENTS

This research was funded by a Canada Excellence Research Chair (CERC) grant to author A.M.O.

## SUPPLEMENTARY FIGURES

**Figure S1.**
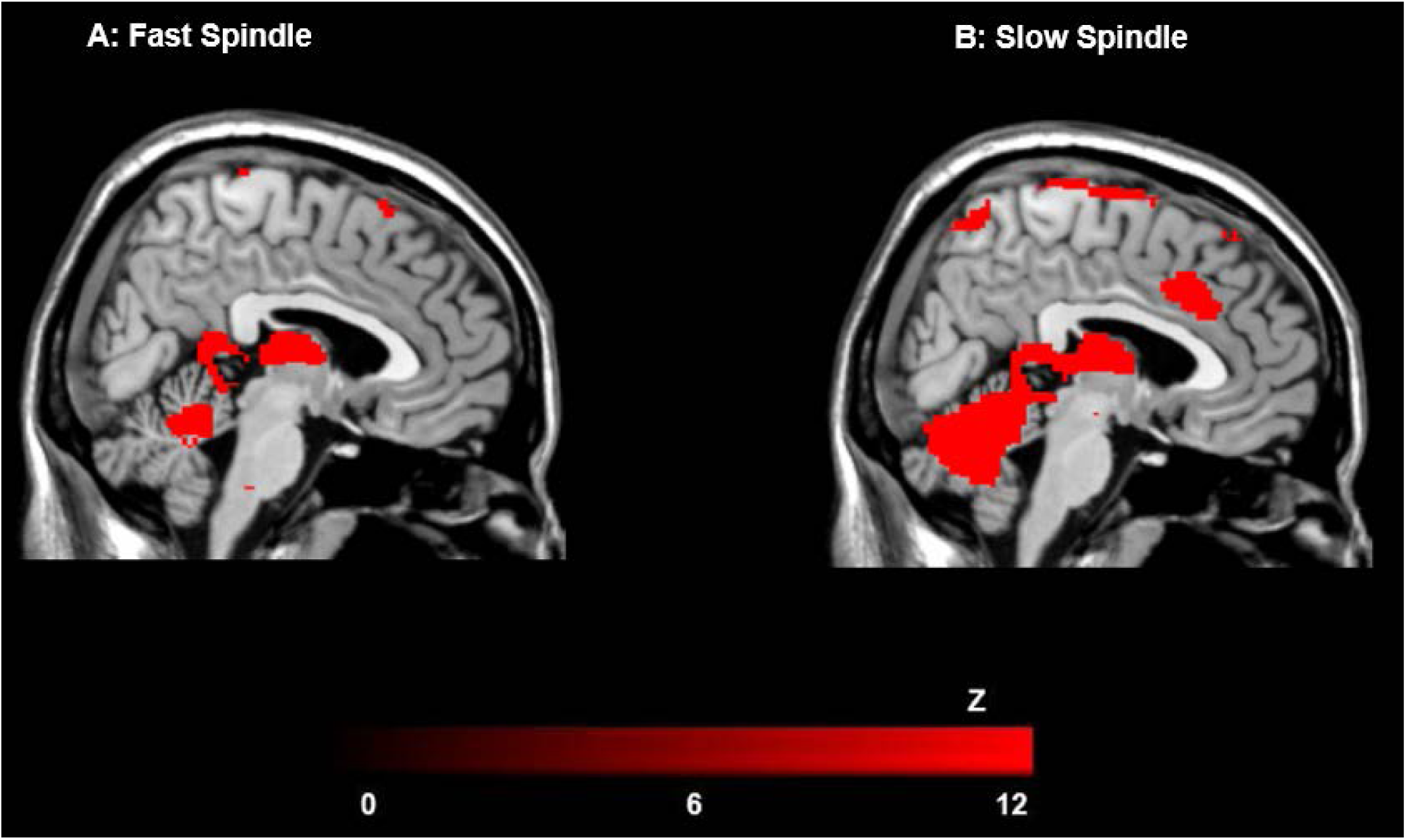
Cerebral activation of fast and slow spindles. Activations time-locked to fast spindles (13.5-16 Hz) at Pz (**A**) and slow spindles (11-13.5 Hz) at Fz (**B**) were similar in all brain areas, but visibly to a greater extent in fast spindles (with the exception of medial frontal activation in slow but not fast spindles). However, there were no significant difference between fast spindles vs. slow spindle-related activations.

**Figure S2.**
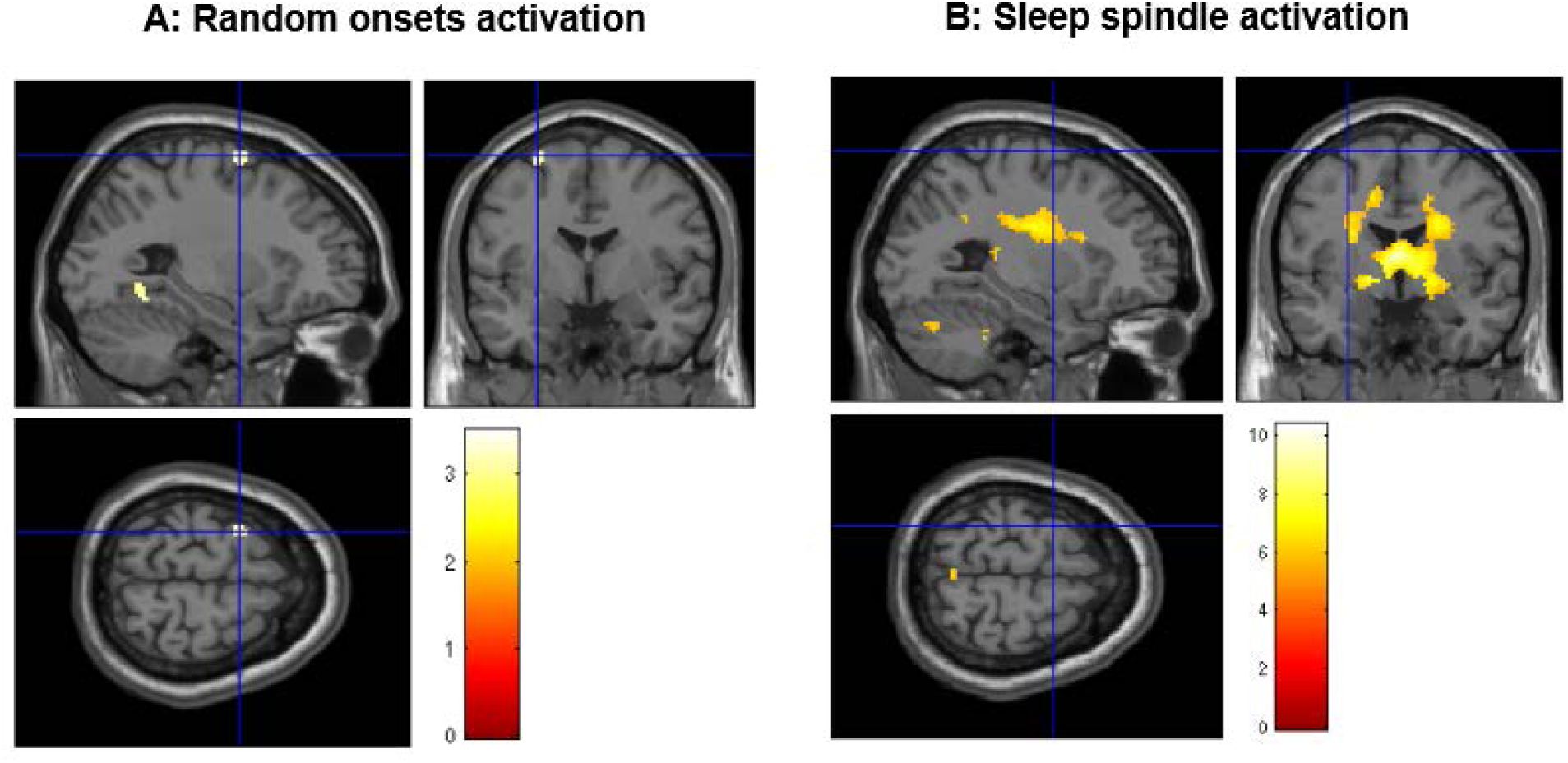
Cerebral activations time-locked to random onsets in NREM stage sleep. The results revealed a small single cluster at left frontal lobe which did not survive FWE correction (peak coordinate: -28, -2, 68; uncorrected p < 0.005) (**A**), and did not overlap with activations time-locked to spindles (**B**).

